# Cheats can boost the success of a cooperative invader

**DOI:** 10.1101/2025.04.30.651436

**Authors:** Luke Lear, Angus Buckling, Elze Hesse

**Author notes:** Corresponding author;. Address: Environment and Sustainability Institute, University of Exeter, Penryn, Cornwall, United Kingdom, TR10 9FE. Invasion success, cooperation, *Pseudomonas aeruginosa*, siderophores, diversity, population fitness.

## Abstract

Successful biological invasions are dependent on the invader being able to grow and reproduce in the new environment. One way that microbial invaders may facilitate this is to use cooperative public goods, such as metal-binding siderophores. However, siderophore production can be exploited by non-producing cheats who benefit from production without paying any associated costs. Here, we test the importance of cooperation for the success of *Pseudomonas aeruginosa* invading a 5-species microbial community. We do this by comparing the success of a siderophore-producing strain, a siderophore-deficient mutant strain and a 50:50 mixed population, both in environments with weak (copper absent) and strong (copper present) siderophore requirement. We found no effect of invader type on success when siderophores were less essential for growth, but large differences when they were selectively favoured. Here the producer-cheat mix had the greatest success, with both strains having near equal fitness and reaching high densities, whilst in isolation producers had intermediate success and cheats the lowest. Similarly, resident diversity only differed across invader treatments when copper was present. In conclusion, we show that the presence of cheats can provide a larger benefit for invasion success than pure cooperator populations, but only when public goods are particularly beneficial.

## Introduction

Invasions by microbes can significantly shape the diversity and function of resident microbial communities (1, 2). Although often detrimental, such as in the case of reduced ecosystem function (3) or pathogen introduction (4–6), invasions can also be beneficial, for example in the case of restoring ecosystem function, biocontrol agents (7, 8) or probiotics (9–12). As such, understanding traits that may affect an invading population’s success is of high importance.

Invaders need to be able to grow and reproduce in new environments in order to successfully colonise them (13, 14). This process is commonly restricted by strong competition for space and resources with the resident community, who may have additional advantages from priority and dominance effects (15, 16). Consequently, traits that offer a competitive advantage for the invader are likely to increase its success. Such traits could include those directly associated with acquiring resources (e.g. excreting enzymes), those associated with restricting the growth of competitors (e.g. excreting antimicrobials (17)), or those that change the abiotic conditions to benefit the invader more than the residents (e.g. altering the pH (18) or disturbance regime (19)). The benefits of these traits may be context dependent, offering an advantage in one environment, but not in another. For example, environments facilitating the greatest invader success can be different between two different genotypes of the same bacterial invader (6).

One way in which microbes can increase their fitness is to release extracellular siderophores into the environment (20, 21). These compounds can be used to scavenge essential nutrients and/or to improve growth conditions by detoxifying toxic metals. For example, *Pseudomonas aeruginosa* releases the siderophore pyoverdine in order to both scavenge iron and, when present at high concentrations, detoxify harmful metals such as copper, zinc, and nickel (22). This can be highly beneficial, with mutant strains deficient in pyoverdine production having reduced fitness in both iron-limited (21) and toxic-metal conditions (23).

As siderophores are extracellular, metabolically costly to produce, and can benefit more than just the producer cell, they can act as a public good (24). Consequently, this cooperative trait is subject to invasion by social ‘cheats’ who benefit from the siderophores without paying the cost of production (24). This can ultimately lead to coexistence of producers and cheats, with the relative fitness of either genotype being dependent on both their frequency and the environment. Cheat fitness increases when the benefits of siderophores are high and their production is upregulated, i.e. when either iron is limited or the concentration of toxic metals is high (23). The increased cost under these conditions is because genes associated with siderophore production are induced and production is upregulated. Consequently, this can result in population-level production being reduced when siderophores are of the greatest benefit (23). Whether producers and cheats can be stably maintained in a population is affected by factors such as population density (25), resource abundance (26), and spatial structure (27).

Importantly, siderophore-bound iron can only enter a cell that has the specific receptor for that siderophore-iron complex (28). Therefore, not all species within the community can benefit from siderophores excreted for iron uptake. This means that by scavenging iron from the environment, producers can lock away iron, and consequently restrict access to only the cells that have the specific receptor (29, 30). In contrast, however, the detoxifying benefits of siderophores are not species specific, with low- or non-siderophore producing species benefiting from the presence of siderophore-producing species in metal-polluted environments (31). This is because unlike siderophore-iron complexes, detoxifying siderophores stop toxic metal ions from passively entering cells where they may cause damage (32). Their function is consequently not reliant on specific membrane receptors and can therefore benefit neighbouring cells. While a siderophore-producing invader should experience a competitive advantage in iron-limited conditions (by locking away iron), such invaders could be outcompeted when other species exploit the detoxifying benefits of siderophores in metal-polluted environments. However, it is noteworthy that whilst detoxifying siderophores can benefit non-producers, they are still likely to provide the biggest benefit to the producers, and related kin, as they will be in closer proximity to the area detoxified by the siderophores, especially in spatially structured environments (33). This is supported by our previous work: while producers can provide protective benefits to other species in metal polluted environments, this does not result in siderophore-exploitation (31, 33). Moreover, metal-pollution shifts microbial communities towards siderophore-producing genera (34), demonstrating that producers can have a strong selective and competitive advantage. Furthermore, as siderophores have a greater affinity for iron than for other metals (32), siderophores released to detoxify toxic metals could potentially lock away iron before they bind to the toxic metals.

Populations comprised of just cooperators (producers) are typically thought to have greater fitness than those containing cheats (e.g. ‘conspiracy of the doves’ (35)). For example, populations of siderophore-producing *Pseudomonas aeruginosa* can outcompete *Burkholderia cenocepacia* under iron-limited conditions, but only in the absence of cheats (21). However, other studies have found that populations comprised of both producers and cheats display greater fitness than producer-only populations in yeast (36). The main reason proposed for the latter is that when there are more cheats in the population, there are relatively fewer resources being created by the producers (e.g. available iron), thereby reducing growth and increasing overall resource use efficiency (36). In either case, it is likely that diverse social traits will cause differences in population dynamics and invader success.

Here, we experimentally test the importance of siderophore-based cooperation in determining invasion success, and how dependent this is on the abiotic environment. We do this by testing whether the invasion success of the opportunistic bacterial pathogen *P. aeruginosa* differs when its population is solely comprised of a siderophore-producing strain or siderophore-deficient mutant, or when two genotypes co-invade at a 50:50 ratio. We test this in two environments that are not iron limited: one that contains copper at a concentration where siderophores are selected for as they are beneficial for growth, and one without copper where there is less requirement for siderophores as they provide little benefit. These environments should impose stronger and weaker selection for cheats, respectively (23), and consequently impact the relative fitness of the two invader types (21). We focus on the detoxifying role of siderophores, rather than iron acquisition, because in principle siderophores could benefit or hinder invasion in this context as they can be a more community-wide public good. As well as invader success, we also test whether invader type impacts resident community diversity and composition.

## Methods

### Bacterial strains

The model resident community and invader used were the same as in our previous work (37), with the exception that a siderophore-deficient invader genotype was also used. The resident community is comprised of five bacterial species, isolated from a single compost sample, that can stably coexist in diluted tryptic soy broth (TSB) media for at least 60 weeks (38). The five members are: *Achromobacter sp*., *Ochrobactrum sp*., *Pseudomonas sp*., *Stenotrophomonas sp*., and *Variovorax sp*.. For our invader, we used two genotypes of the opportunistic bacterial pathogen *Pseudomonas aeruginosa* PA01: the wildtype as our cooperative producer genotype, and an isogenic siderophore-deficient mutant strain of PA01 (PAO1ΔpvdDΔpchEF (39)) that cannot produce pyoverdine, or the secondary siderophore pyochelin, as our cheat. *P. aeruginosa* can successfully invade a range of environments, and the producer genotype used here has previously been found to have treatment-dependent success and impact on composition when invading the resident community (37, 40). Crucially, all 7 strains (including both *P. aeruginosa* genotypes) have visually distinct colony morphotypes when plated onto King’s medium B (KB) agar.

To create our resident communities, we first grew each of the five resident species in monoculture for 48 hours at 28°C in static glass vials (microcosms) containing 6mL of diluted (1/64^th^) TSB. Equal volumes of the monocultures were then combined to form a mixture of the five species, of which 60μL was added to 8 replicate microcosms containing 6mL of diluted TSB. These were then kept static at 28°C with loose lids to allow oxygen transfer – the same conditions used throughout. After eight days, each of the eight communities was both plated onto KB agar to quantify its density, and transferred into eight different microcosms, yielding a total sample size of 64 microcosms. Half of these contained diluted TSB and half contained diluted TSB supplemented with copper sulphate (CuSO_4_) at a final concentration of 50µM.

The two invader genotypes were grown statically for 24 hours in 6mL of KB at 28°C. We then adjusted their optical density to 0.75 at 600nm before diluting these cultures 10-fold in M9 salt buffer. This resulted in invader doses of 1.51 × 10^6^ producer CFU and 1.20 × 10^6^ cheat CFU: equal volumes of these cultures were mixed to create the 50:50 mix treatment (actual proportion = 44.2% cheats). As the mean density ± SD of the resident in the fresh microcosms was 1.43 x 10^7^ ± 3.14 x 10^6^ CFU, this resulted in an initial invader proportion of around 10%.

### Pyoverdine assay on the ancestral cooperative invader strain after growth with and without copper

Previous work on copper-induced siderophore production in *P. aeruginosa* has used higher copper concentrations and different media to those used here (23). We therefore tested if our two environments resulted in different densities of the two invader types when they were grown in monoculture and whether they resulted in different pyoverdine production by our producer strain. To do this, 16 microcosms were inoculated with monocultures of the producer strain of *P. aeruginosa* and 16 with monocultures of the cheat strain as described above; 8 of the 16 microcosms contained diluted TSB and 8 contained diluted TSB supplemented with copper sulphate (CuSO_4_) at a final concentration of 50µM. On day 4 microcosms were homogenised before 20µL of culture was diluted and plated on to KB agar and 20µL was transferred into three wells of a 96-well plate containing 180µL of KB media made iron limited through the addition of a strong iron chelator (200 µM 2,2’-bipyridine and 25 mM HEPES buffer (20)) and incubated for a further 48 hours at 28°C. This iron-limited common garden environment stimulates siderophore production. Using a Varioskan LUX microplate reader the plate was then shaken and pyoverdine quantified by measuring the optical density at 600 nm (OD_600_) and the fluorescence at 460 nm following excitation at 400 nm of each culture (41). The OD_600_ measurements were used to estimate pyoverdine production per capita using fluorescence (relative fluorescence units: RFU) / OD_600_. The average of the three technical replicates was used in the analysis.

### Experimental design

Our experiment comprised of a factorial design with four invader treatments and two environment treatments, for a total of eight treatments that were each replicated eight times. The four invader treatments were: producer genotype, cheat genotype, a 50:50 producer-cheat mix, and a non-invaded community-only control; the two environment treatments were the presence or absence of copper. Invaders were added post-transfer on day 8, and the communities left for a further 8 days until they were destructively sampled and frozen in glycerol at a final concentration of 25% and stored at -70°C. Defrosted samples were diluted to a dilution of 10^-5^ in M9 salt buffer, plated onto KB agar, and incubated at 28°C for 48 hours.

### Statistical analysis

#### Model comparison and residual behaviour

All analysis were done in R version 4.1.0 (42). We used the ‘*lme4*’ package (43) for our linear and generalised linear mixed effects models, and the ‘*DHARMa*’ package to check the residual behaviour of our models. A random intercept was included for each community to control for any variation between the communities created in the initial eight days of the experiment before the invader was added. To arrive at the most parsimonious model, we sequentially removed model terms, such as the interaction between invader type and copper, and compared model fits using χ^2^ tests. We then used the ‘*emmeans*’ package (44) to calculate contrasts, with p values adjusted for multiple testing using the fdr method where appropriate. We calculated all possible contrasts for any model with a significant interaction between invader-type and copper. To reduce the effect of adding a single colony when log-transforming density estimates, we use CFU counts per plate (CFU/plate) as response variable instead of CFU/mL, as all cultures were plated at same 10^-5^ dilution.

#### Invader monocultures in the two environments

To test if the copper-polluted and unpolluted environments differentially affected the density of the different invader types in monoculture, we used a linear model with CFU per plate as the response variable and invader type (producer or cheat), environment (copper present or absent), plus their interaction as the explanatory variable. CFU was log(CFU+1) transformed to improve model fit.

#### Pyoverdine production

Per capita pyoverdine production by *Pseudomonas aeruginosa* was compared between the two environments using a 1-way Anova with per capita pyoverdine production as the response variable and copper (presence/absence) as the explanatory variable.

#### Invader success

We used relative invader fitness (v) as our metric for invasion success, as this accounts any differences in resident density across the communities at the time of invasion. This metric is the proportional change of invader compared to the total resident community: v = x_2_(1 − x_1_)/x_1_(1 − x_2_), where x_1_ is the initial invader proportion and x_2_ the final (45), and has been used previously to quantify invader success (6). To test treatment effects on invader success, we used a linear mixed effects model with invader type (factor with three levels), copper (factor with two levels), and their interaction as explanatory variables. To improve model behaviour, v was transformed using log(v+1).

#### Cheat fitness

In order to quantify the fitness of the cheat relative to the producer in the 50:50 mix invader treatments, we used *v* as above, with x1 now being the initial proportion of cheat in the invading population and x2 the final. We then tested whether this was different between the copper treatments using a linear mixed effects model with copper as a factor with two levels as an explanatory variable. In this model cheat fitness (v) was square root transformed to improve model fit. Then we used two separate one sample T tests to test whether the final proportion of cheats in the invader population was different to the initial actual proportion (0.442) in both the presence and absence of copper. Next, we tested whether cheat fitness and population-level invader success were associated using a linear mixed effects model with invader success (log(v+1)) as the response variable and cheat fitness (non-transformed), copper, plus their interaction as the explanatory variables. Finally, we tested whether the cheat had a higher density in monoculture or co-culture with the producer in both the copper-free and copper-polluted environments. To do this we used a linear mixed effects model with log(cheat CFU +1) as the response variable, and both invader monoculture/co-culture and copper present/absent as the explanatory variables.

#### Resident and invader density

To determine treatment effects on the density of the resident community we used a linear mixed effects model with total number of resident CFU as the response variable, and invader type (factor with four levels), copper (factor with two levels), and their interaction as explanatory variables. Resident CFU was log10+1 transformed to improve model fit. We used a linear mixed effects model with the same explanatory variables to determine treatment effects on the density of the invader (this was also log10+1 transformed to improve model fit).

#### Resident diversity

To quantify resident biodiversity, we used the Simpson’s index 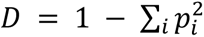 where *p*_*i*_ is the proportion of the *i^th^* species of the resident community (46). We then used a linear mixed effects model with invader type (factor with four levels), copper (factor with two levels), and their interaction as explanatory variables to test if this differed between treatments. To test if resident diversity was associated with cheat fitness, we used a linear mixed effects model with Simpson’s index as the response variable and cheat fitness, copper, plus their interaction as the explanatory variable. Cheat fitness was transformed to log(v+1) to improve model fit.

#### Species densities

We determined treatment effects on the density of each of the five resident species using a generalised linear latent variable model from the *‘gllvm’* R package (47). Taking this multivariate approach allows us to distinguish between treatment effects on densities and the latent interactions among the resident species (i.e. if a species is directly affected by copper or by the effect of copper on a different species). We included invader type, copper, and their interaction as the environmental variables, and used the non-invaded, copper-free treatment as the reference level.

## Results

### Siderophore production was selected for, and beneficial in, the copper environment

Here, we tested the importance of cooperation for the success of a microbial invader, and whether this was dependent on the environment being invaded. We first determined whether our copper environment selected for greater siderophore (pyoverdine) production by *P. aeruginosa* when it was grown in monoculture. We found pyoverdine production to increase significantly by 19.9% in the copper-polluted compared to copper-free environment (mean RFU/OD_600_ ± SD in the copper environment = 1107.6 ± 31.6 and in the copper absent environment = 923.9 ± 42.2; F_1,14_=97.0, p < 0.001; Fig. 2A). Next, we tested whether pyoverdine production is beneficial for growth in the copper environment by comparing the density of the producer and cheat invader types when grown in monoculture in the two environments. Copper significantly decreased the density of both invader types (copper main effect: F_1,28_=227.9, p < 0.001; Fig. 2B). However, this effect was different between the two types (copper-invader type interaction: F_1,28_=14.4, p < 0.001; Fig. 2A), with densities being equal across invader types in copper-free media (mean CFU ± SD = 642.5 ± 158.6 for the producer and 772.5 ± 323.7 for the cheat; p=0.28) but the cheat having a significant 44.7% lower density than the producer in copper-polluted environment (mean CFU ± SD = 203.3 ± 39.2 for the producer and 112.5 ± 29.8; p< 0.001). These results demonstrate that copper selects for increased pyoverdine production and that this is beneficial for growth in the copper conditions. That the cheat and producer had statistically equal densities in the copper-free environment implies that pyoverdine is not beneficial for growth in this environment, suggesting that it is not iron limited after four days.

**Figure 1.**
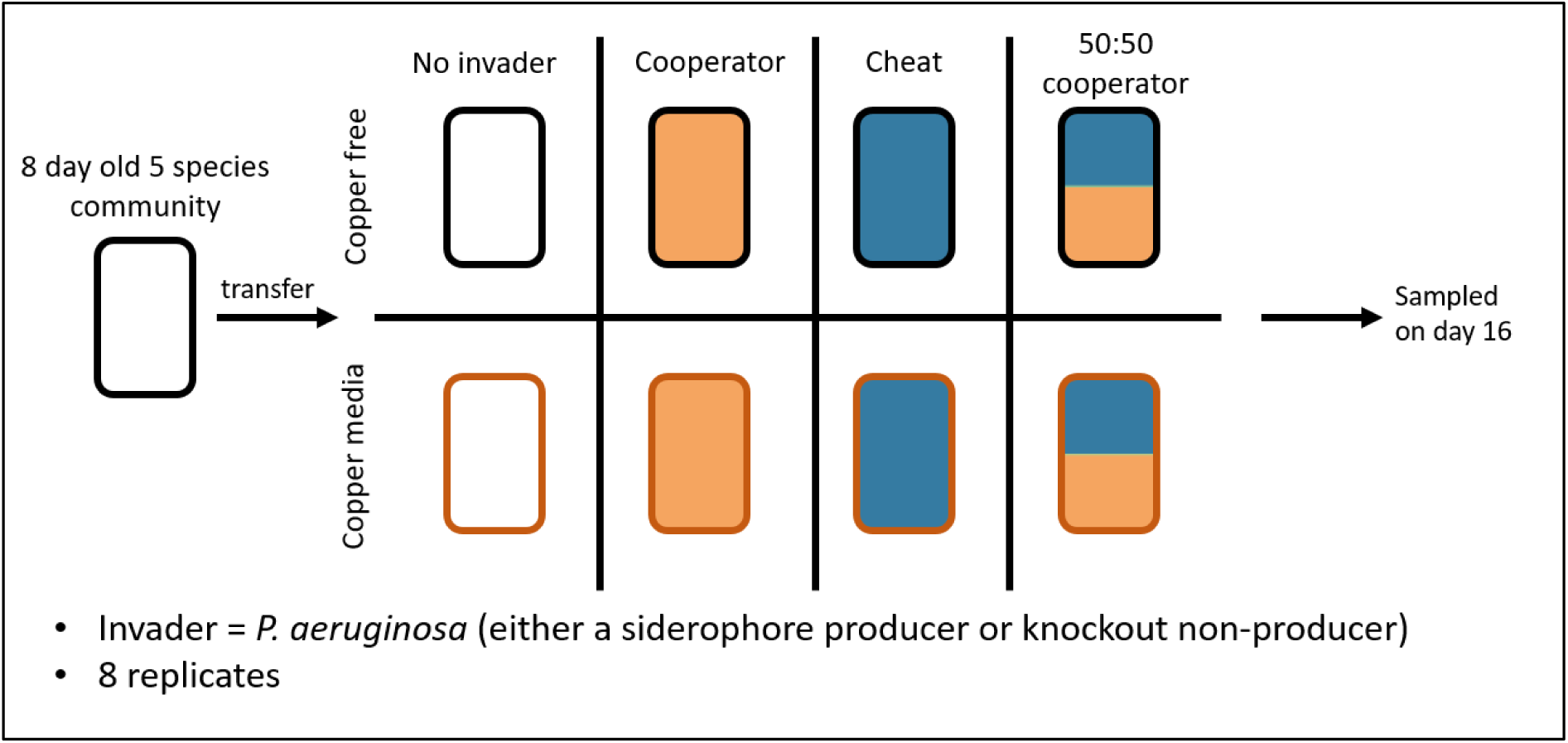
Schematic design used to test the importance of cooperation for the success of an invader. A community of five bacterial species was grown for eight days before being split into one of four invasion treatments (invaded with a siderophore producer, non-producer, 50:50 mix, or no invader) and two environment treatments (copper present or absent), each replicated eight times for a total of 64 microcosms. Samples were taken on day 16 by fully homogenising the cultures and freezing them.

**Figure 2.**
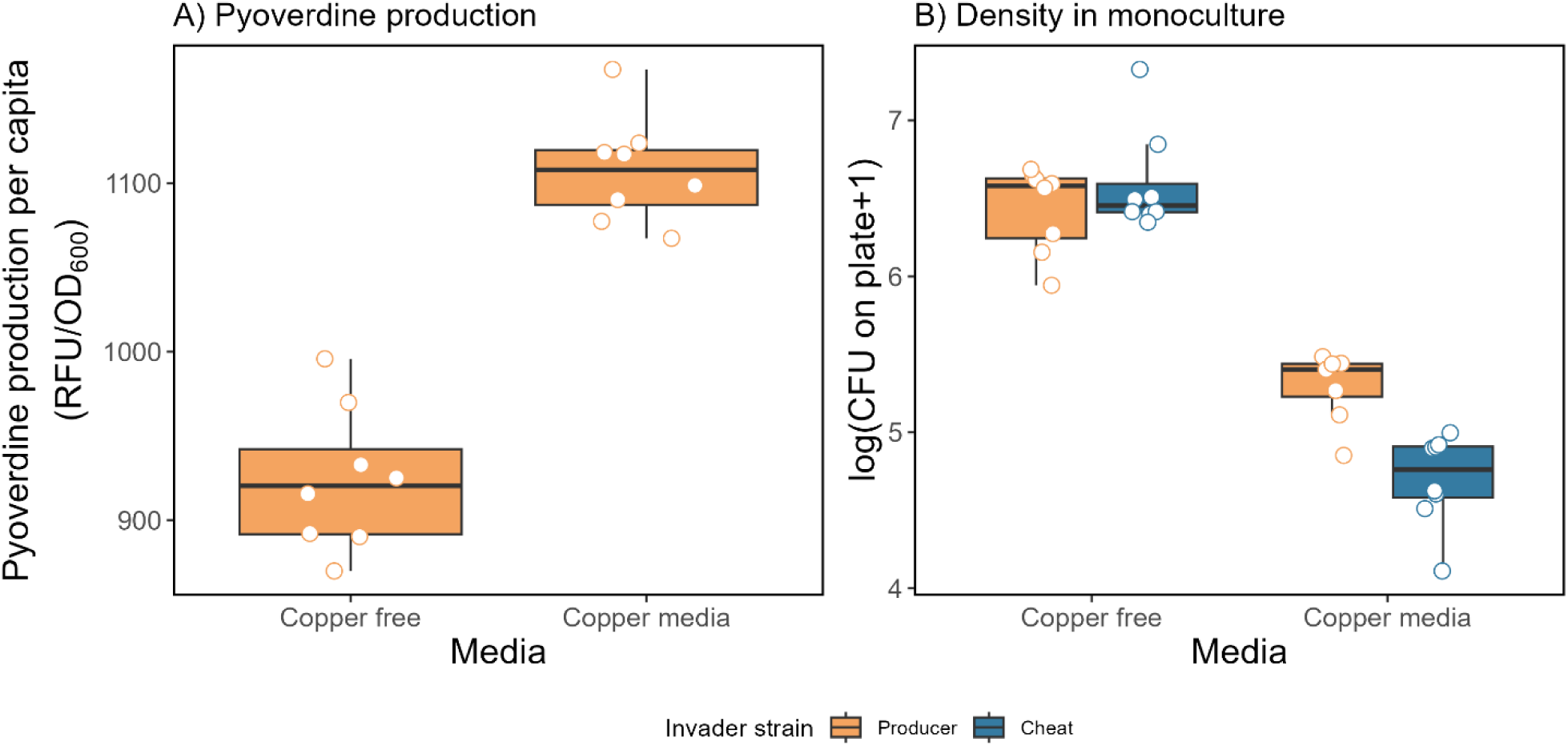
Panel A shows pyoverdine production per capita (standardised relative fluorescence units per OD_600_) by the producer strain of *Pseudomonas aeruginosa* after monoculture growth in diluted TSB or diluted TSB supplemented with copper sulphate (CuSO_4_) at a final concentration of 50µM for 4 days and then for 48 hours in iron-limited KB. Panel B shows the density (log(CFU per plate+1)) of the producer (orange boxes) and cheat (blue boxes) strains of *P. aeruginosa* after 4 days growth in either diluted TSB or diluted TSB supplemented with 50µM CuSO_4_. Circles show individual replicates (n = 8).

### Invader type affected success in one environment, but not in the other

The invader successfully established (had a higher final proportion than initial proportion, i.e. v was > 1) in 47 out of 48 invaded communities. We found invader success to vary across copper environments, and this effect varied across invader types (producer, cheat, or a 50:50 mix), with invader type significantly affecting success in the copper media but not in the copper-free media (invader-copper interaction on invader success: *χ^2^* = 39.2, d.f. = 2, p < 0.001; Fig. 3). In the copper-free media, the three invader types did not statistically differ in their success (p=>0.054 for all contrasts) and had a mean success (log(v+1) ± SD) of 3.17 ± 0.57. However, in the presence of copper, invader types differed significantly from one another in terms of their invasion success, with the cheat’s success (mean log(v+1 ± SD) = 1.44 ± 1.04) being significantly lower than both the producer’s success (mean log(v+1 ± SD) = 2.78 ± 0.96; p<0.001) and the mixed invader’s success (mean log(v+1 ± SD) = 5.15 ± 0.91; p<0.001), with the mixed being higher than the producer’s (p<0.001). Interestingly, we found the producer’s success in copper to be statistically equal to its success in the copper-free medium (p=0.35), whereas copper reduced the success of the cheat invader by 66% (p<0.001), having a large positive effect on the mixed invader, increasing its success by 627% (p<0.001). These results therefore demonstrate that a mix of cooperators and cheats can lead to the greatest invasion success, but only when there is higher requirement for the public good (the production of siderophores to detoxify copper).

**Figure 3.**
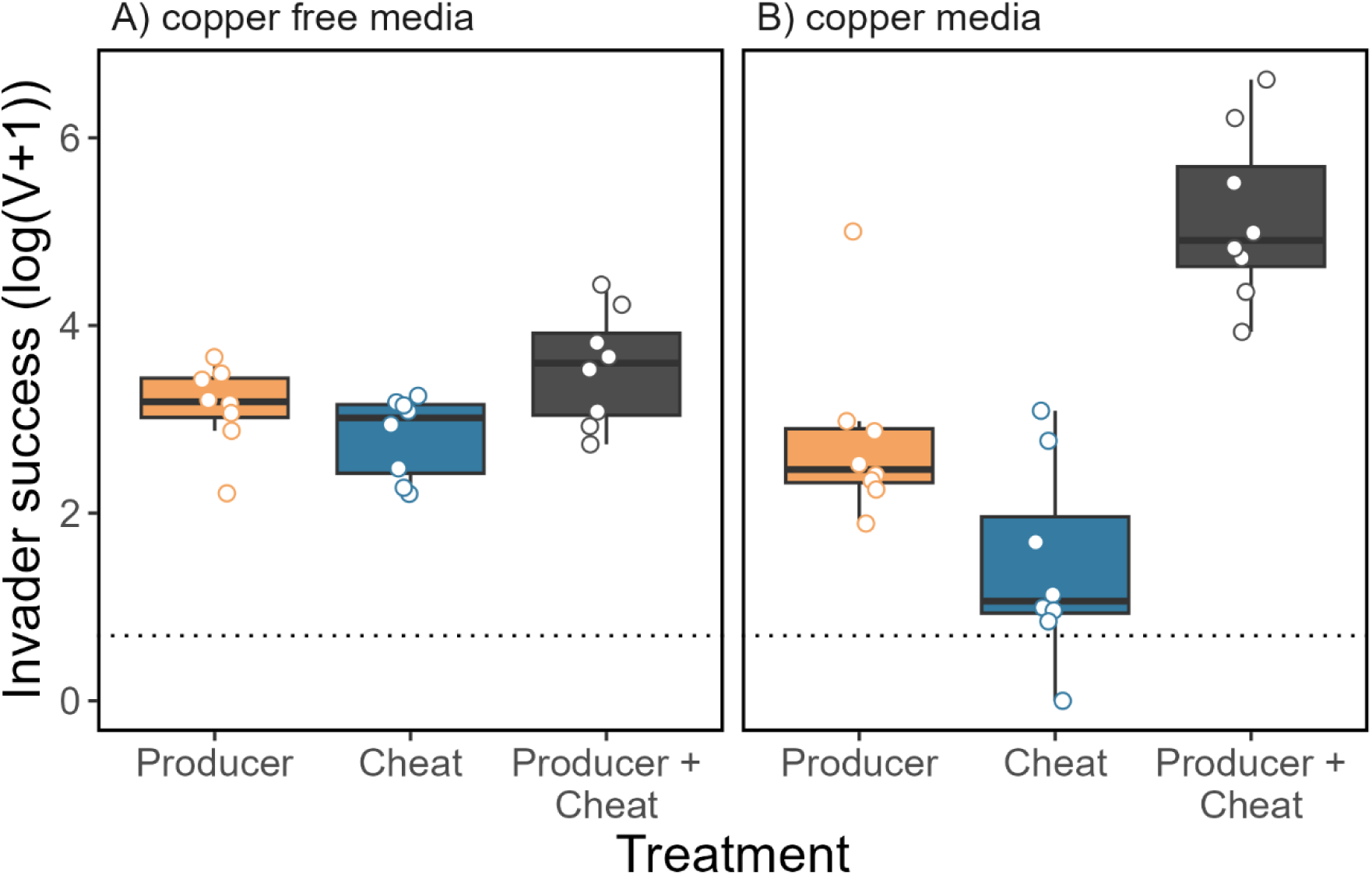
The success (log(v+1), where v is the temporal proportional change of the invader when invading a resident 5-species community. Invader populations either consisted of a siderophore-producing genotype (‘Producer’), a non-producing genotype (‘Cheat’), or a 50:50 mix of the two genotypes (‘Producer + Cheat’). Resident communities were grown in copper-free (Panel A), or copper-polluted environments (Panel B). The horizontal dashed line at 0.69 (log(1+1)) shows the threshold for successful invasion: values above this demonstrate the invader population has increased in frequency over time. Circles show individual replicates (n = 8).

### Cheats benefit from producers only when copper is present

To determine why having a mix of producers and cheats was beneficial for invading a microbial community in the presence, but not absence, of copper we compared the final relative fitness of the cheat to that of the producer in the mixed-invader treatment across copper environments. We found copper to positively affect cheat fitness (copper main effect: *χ^2^* = 55.5, d.f. = 1, p < 0.001; Fig. 4A), with the cheat always being outcompeted when copper was absent (mean cheat fitness without copper ± SD = 0.047 ± SD 0.047), but having a fitness advantage in the presence of copper (mean cheat fitness with copper ± SD = 2.38 ± 1.82; Fig. 4A). Consequently, the cheat comprised on average 57.3% of the invader population in the presence of copper, but only 0.04% in the absence. Comparing this to the initial invader proportion (44.2%), we find the that the slight increase in final cheat proportion in the presence of copper was not significant (p=0.10; Fig. 4C), but that the decrease in the absence of copper was (p<0.001; Fig. 4D).

**Figure 4.**
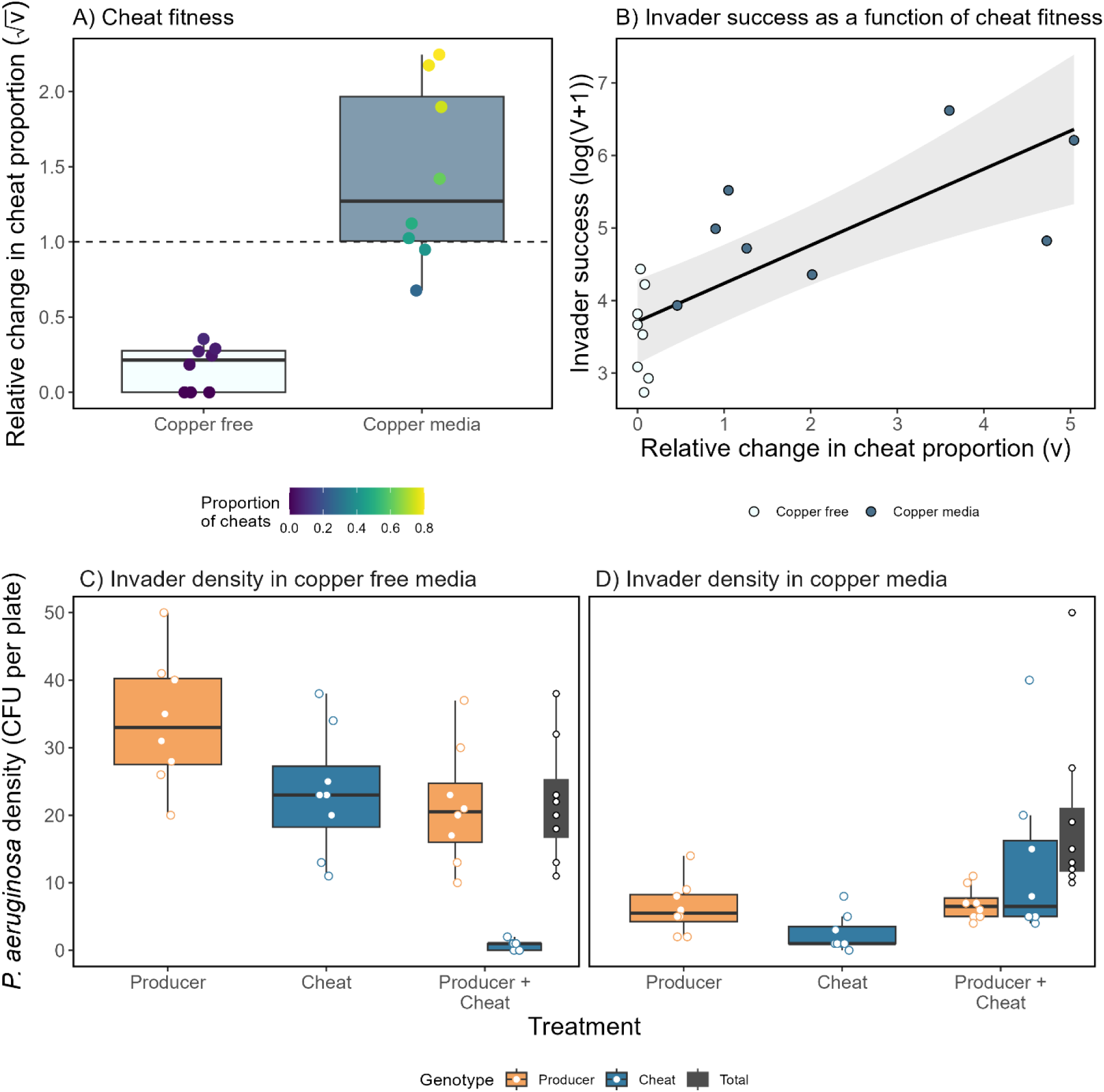
Relative cheat fitness (measured as proportional change in the ratio of cheat versus producer genotype) when (A) grown in co-culture with the producer in the two copper treatments and (B) its effect on population-level invader success. Panel C shows the CFU per plate of the two invader genotypes when invaded into copper-free medium and panel D when invaded into copper-polluted medium. Circles show individual replicates; in panel A the colours show the proportion of cheats in the population (purple = low values, yellow = high), in panel B the copper treatment (orange = absent, blue = present), and panels C and D the invader genotype (orange = producer, blue = cheat, and black = total). The line in panel B shows the best model fit and the shaded area the 95% confidence interval.

We then tested whether increased cheat fitness in the mixed invader populations was associated with greater overall invader success, as it would allow an overall greater invader population density, and found it to be true (cheat fitness main effect: *χ^2^* = 5.01, d.f. = 1, p= 0.025; Fig. 4B). Furthermore, we found this effect to be statistically equal between the two environments (cheat fitness-copper interaction: *χ^2^* = 0.48, d.f. = 1, p= 0.49).

Finally, to determine whether the cheat benefited from and exploited the producer (i.e. whether siderophores acted as an intraspecific public good), we tested whether the cheat density was greater when the producer was present compared to when it invaded as monoculture. We found that in the absence of copper, the cheat reached a density > 30 times greater in monoculture (mean CFU ± SD = 23.4 ± 9.27) than in the presence of the producer (mean CFU ± SD = 0.75 ± 0.71) (p < 0.001), demonstrating that intraspecific competition between the two genotypes was strong and that the producer was fitter (Fig.4A&C). However in the presence of copper, the cheat significantly benefited from the producer and reached a density > 5 times greater in co-culture (mean CFU ± SD = 12.75 ± 12.42) compared to in monoculture (mean CFU ± SD = 2.50 ± 2.73; p=<0.001; Fig. 4D), demonstrating that the cheat strongly benefited from the producer in the presence of copper where pyoverdine production was greater but not in the absence. Overall, these results demonstrate that in the presence of copper the cheat has increased fitness through benefiting from the producer and can consequently maintain its population size. Interestingly however, the increase in cheat fitness had no negative impact on the density of the producer (suggesting that it was not exploited) and, as a consequence, total invader density of the mixture was higher than in monoculture in copper-polluted environments.

### Copper reduced both resident and invader density

To determine the effect of invasion on resident density, we tested treatment effects on total resident density (Fig. 5). The copper treatment significantly reduced the density of the residents from 119 ± SD 58.8 to 32 ± SD 22.2 CFU per plate (copper main effect: *χ^2^* = 139.0, d.f. = 1, p < 0.001), However, the severity of the reduction depended on the invader treatment (invader-copper interaction: *χ^2^* = 9.46, d.f. = 3, p = 0.024). In the presence of copper, we found the cheat to decrease resident density less than both the producer (p=0.01) and the mixed invader (p=0.002), but not to differ to the non-invaded control (p=0.17). The producer, mixed, and non-invaded control all did not differ in their resident densities (p=>0.23 for all contrasts). We note that when copper was absent, both the producer and cheat invaders significantly reduced resident density compared to the control (p = 0.008 & 0.034, respectively). The same trend occurred for the mixed invader, but this was not significant (p=0.051). The three invader types reduced the resident density equally (p=>0.47 for all contrasts). Finally, the copper treatment also significantly reduced the density of the invaders (*χ^2^* = 83.0, d.f. = 1, p< 0.001), so that whilst invader density was lower in the copper treatments (Fig.3D), this is not a reflection of its success.

**Figure 5.**
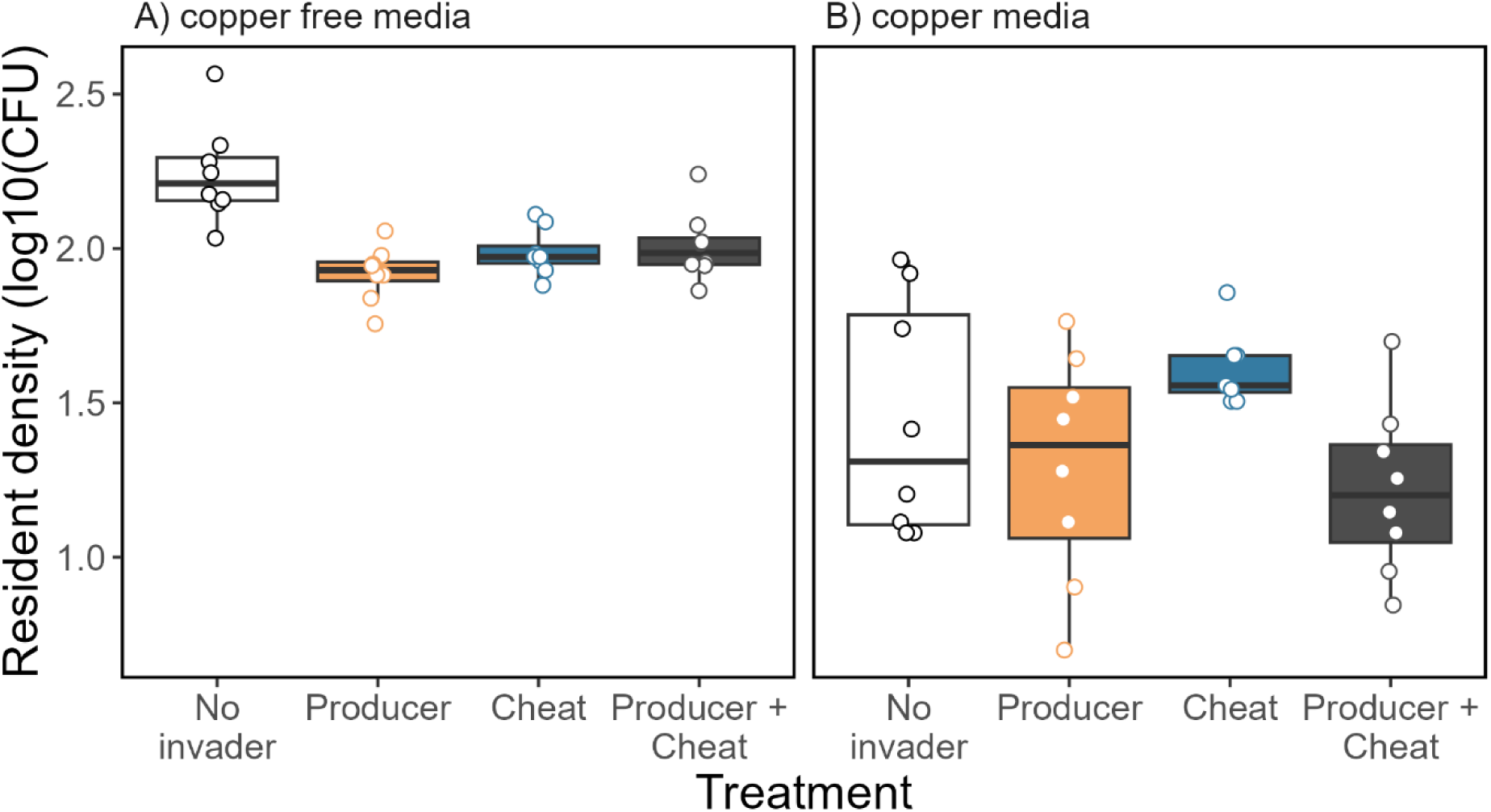
The density (log10(CFU per plate)) of a resident community after being invaded by *Pseudomonas aeruginosa* populations that were either siderophore producers (orange), non-producers (blue), or a mix (grey); a non-invaded control was also used (white). Invasions either took place in (A) a copper-free environment or (B) an environment containing 50µM of copper. The circles show individual replicates (n=8).

### Invader types have differential effect on resident diversity

Resident diversity (Simpson’s index) was significantly affected by invader type, depending on the copper environment (invader-copper interaction: *χ^2^* = 11.0, d.f. = 3, p=0.016; Fig. 6A). In the copper-free environment there was no significant difference in diversity between any of the invader treatments, including the non-invaded control (mean diversity in copper-free media ± SD = 0.56 ± 0.09; p=>0.056 for all contrasts; Fig. 6A). However, once again we found differences across invader types in the copper medium (Fig. 6B), with invasion by the cheat monoculture significantly reducing diversity to a mean Simpson’s index of 0.34 ± 0.10 SD compared to that of 0.48 ± 0.09 SD in the non-invaded control, and 0.50 ± 0.14 SD when monocultures of the producer were added (p=0.046 & p=0.016, respectively). Communities invaded with mixed populations in copper-polluted conditions had diversity values (0.46 ± 0.17 SD) between those of the cheat and producer treatments and did not significantly differ to either (p=0.056 & p=0.055, respectively), nor to the non-invaded control (p=0.85). Communities invaded with the producer did not have statistically different diversity to the non-invaded control (p=0.85).

**Figure 6.**
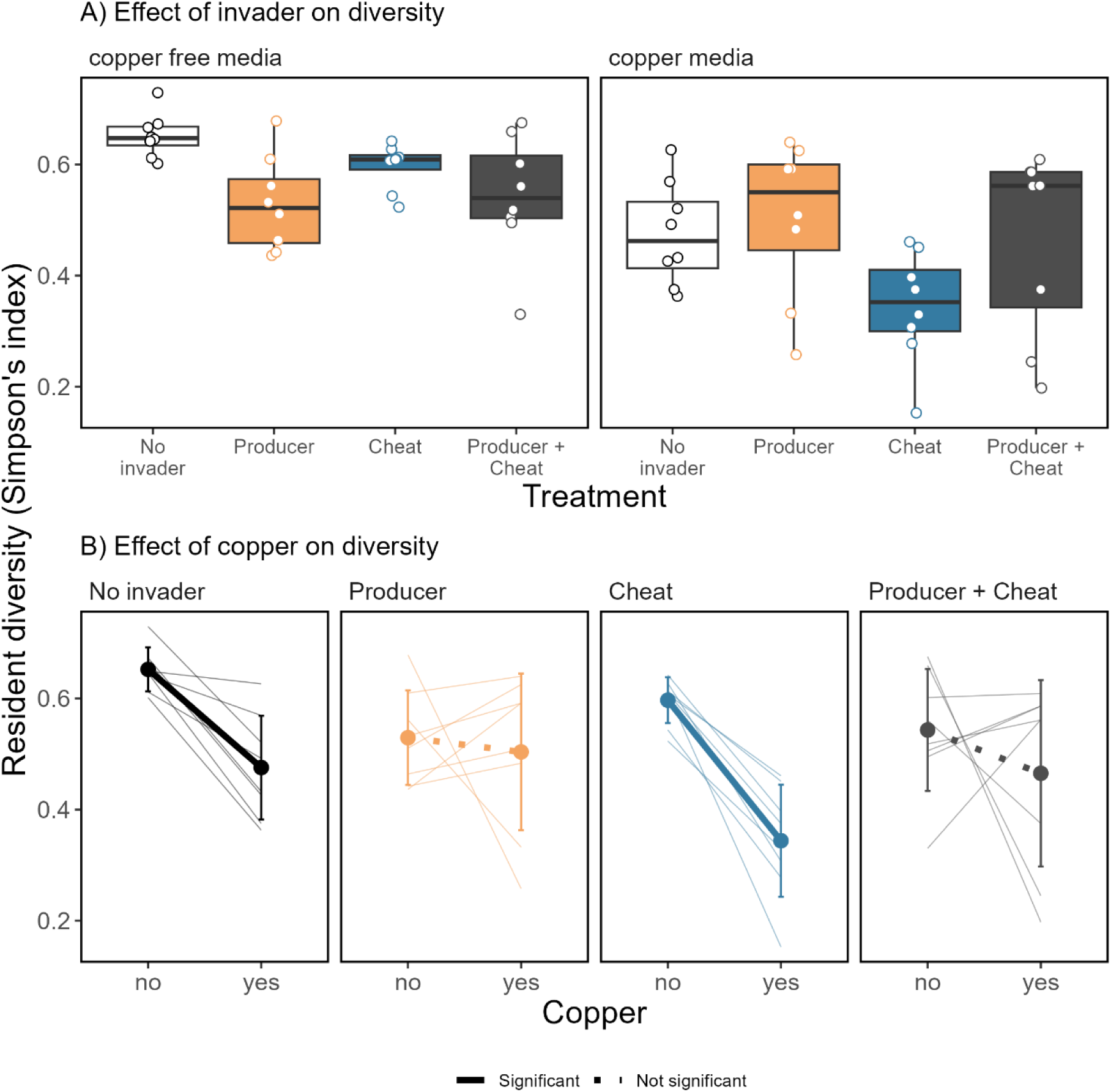
The diversity (Simpson’s index) of a resident community after being invaded by *Pseudomonas aeruginosa* populations that were either siderophore producers (orange colours), non-producers (blue colours), or a 50:50 mix of both (grey colours); a non-invaded control was also used (white/black colours). Invasions either took place in a copper-free environment or an environment containing 50µM of copper. Panel A compares diversity between invader types whereas Panel B compares diversity between the two copper environments, with the large points and thick lines showing the mean of the 8 replicates, the thin lines showing each replicate, and the error bars the standard deviation. In panel A the small circles show individual replicates. Solid thick lines in panel B show a significant difference in diversity between the copper treatments (p=<0.05), whereas dotted lines show non-significant differences.

Interestingly, when a producer invaded, either in monoculture or co-culture with the cheat, resident diversity was equivalent in copper-free and copper-polluted media (p=0.68 & p=0.26, respectively; Fig. 6C). However, when no invader or monocultures of the cheat invaded, diversity was significantly reduced in the copper treatments compared to the copper-free treatment (p=0.007 & p<0.001, respectively). Despite this, we found the fitness of the cheat to not significantly affect resident diversity (*χ^2^* = 0.003, d.f. = 1, p=0.96). This suggests that the presence of a producer mitigates the negative effect of cheats on diversity, even when cheats are highly prevalent.

### Invasion and copper affected the density of each resident species differently

To explore how treatments effected resident diversity, we used a multivariate approach determine how invader type, copper, and their interaction affected the density of each of the five resident species, using the non-invaded, copper-free treatment as our reference. We found copper to have a significant large negative effect on the density of *Pseudomonas*, *Ochrobactrum*, and *Stenotrophomonas* (p=<0.001 for all contrasts), but to not significantly affect the density of *Achromobacter* or *Variovorax* (p = 0.54 and 0.082, respectively; Fig.7). *Pseudomonas* and *Ochrobactrum* were significantly reduced by all invader types (p=<0.013 for all contrasts), this effect was not different between the copper treatments as *Pseudomonas* and *Ochrobactrum* are highly susceptible to copper (i.e. the effect of invasion was masked by the large negative effect of copper). Invasion did not affect *Stenotrophomonas* in the absence of copper, and only the cheat affected the density of *Stenotrophomonas* in the presence of copper: it significantly reduced it (p=0.023), whereas the producer and producer-cheat mix had no significant effect (p= 0.68 and 0.11, respectively). *Achromobacter* was not significantly affected by invasion both in the presence and absence of copper (p => 0.15 for all six contrasts). *Variovorax* was significantly reduced by all three invader types in the absence of copper (p=<0.009 for all contrasts). However, in the presence of copper, it significantly benefited from invasion by the producer (p=<0.001) and the cheat (p<0.001), but not the producer-cheat mix (p=0.075). This benefit resulted in *Variovorax* reaching a higher density in these treatments compared to the non-invaded copper-free treatment, demonstrating that as copper did not affect its density invasion positively affected *Variovorax* density.

**Figure 7.**
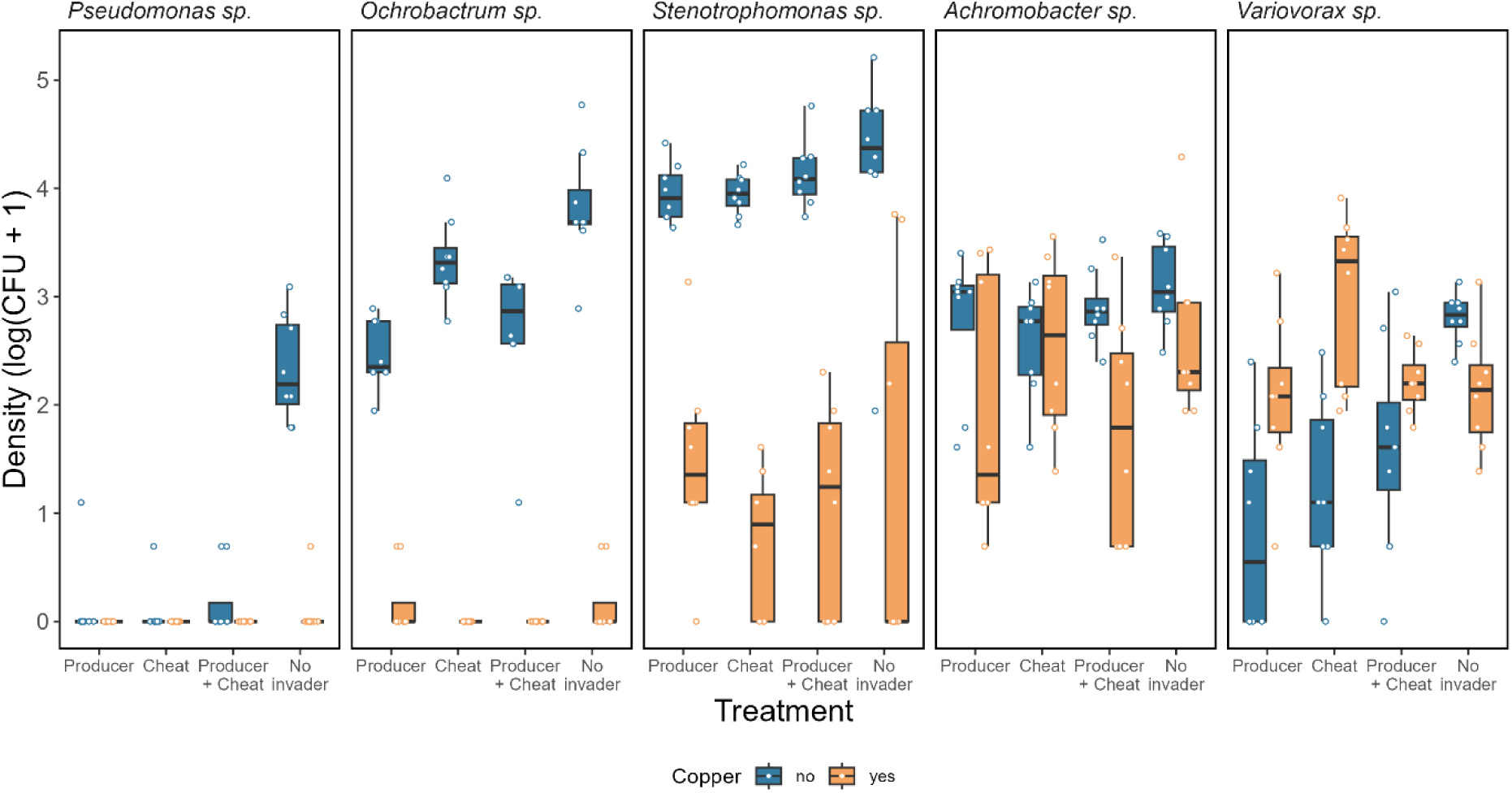
The density of the five resident species after being invaded by *Pseudomonas aeruginosa* either as a siderophore-producing genotype (‘Producer’), a non-producing genotype (‘Cheat’), a 50:50 mix of the two genotypes (‘Producer + Cheat’), or a non-invaded control (‘No invader’). Blue boxes show when they were invaded in a copper-free environment and orange boxes when copper was present. Circles show individual replicates (n=8).

## Discussion

Here, we experimentally tested the importance of cooperation for successful invasion of a microbial community in two different environments: one where the cooperative trait – siderophore production – is of lower benefit (in the absence of copper) and one where it is of greater benefit (in the presence of copper). We found that, consistent with their growth in monoculture, different types of invaders had a similar invasion success in copper-free conditions, where siderophores were less beneficial for growth. However, invader type did significantly affect invasion success in the presence of copper where siderophores were beneficial, with all three invader types having different success. This was because here siderophores were required more, resulting in the cheat having reduced success when invading on its own but the producer being equally successful in presence and absence of copper. However, when the cheat and producer invaded the copper environment in coculture, invasion success was the greatest overall. This was because the increased requirement for siderophores resulted in the cheats benefiting from the producer and consequently in them having similar fitness – the cheat could subsequently persist alongside the producer causing a larger invading population. We therefore show that cooperation can play an important role in invasion ecology, but only when the benefits of the cooperative trait are sufficiently high that the cheat’s fitness is increased. Moreover, we found a similar result with the invader’s impact on the resident composition – invader type was important but only in the copper environment where siderophores were under greater selection and the cheat could persist alongside the producer.

Invasion success was on average equal between the two environments, with producers having similar success across both and, in the copper environment, the greater success of the mixed-invader balancing the reduced success of the cheat. Considering that classical game theory (e.g. conspiracy of the doves) predicts population fitness to be greatest under full cooperation (35), it is perhaps surprising that we did not find monocultures of producers to be the most successful invader in any scenario. Instead, we found the presence of cheats to boost the fitness (density) of the invading population in the copper environment, and cheats to have no effect on success in the absence of copper. Cheats boosting population fitness has previously been found in yeast, where populations that contained producers and non-producers of an extracellular enzyme reached greater densities than either in monoculture (36). In that work, it was suggested that cheats can boost population fitness when resources are used inefficiently, when the amount of cooperation cannot be accurately assessed, and when population structuring results in producers accessing more of the benefit of the public good than the cheat (36). Here, it is most likely that siderophores, produced to detoxify copper, are helping the cheat maintain higher populations densities compared to when it is invading in monoculture. That this does not significantly reduce the fitness of the producer strain could be because the extra-cellular nature of the siderophores mean that the producer and cheat can benefit from their direct protective effects simultaneously (i.e. they do not compete for their benefit). This effect is made more likely by the overall reduction in population density caused by the copper reducing competition between the cheat and producer (31) and altering their relative fitness (25). We therefore show that the impact of cheats on population fitness is dependent on both the presence of other species and the benefit of the public good in question. Importantly, our work demonstrates that these differences in population cheat-producer dynamics can determine the success of microbial invaders.

The fitness of the cheat in our study is overall lower than that in previous work (23), with the presence of the community being the most likely reason for this (48). For example, in our study the relative fitness of the cheat is similar to that of the producer in the copper media, whereas in previous work cheats reach much higher frequencies in copper polluted media (23). Moreover, we found the cheat to be highly selected against in the absence of copper, and to consequently be outcompeted to near extinction by the producer when both were competing with other community members for resources. Whilst the severe extent of this fitness loss is unexpected based on previous work in single-species populations of *P. aeruginosa*, it supports recent findings that siderophore producer-cheat dynamics are different in the presence of a community (48). This is likely due to the resident community also competing with the cheat, for example for iron which may have become limited by eight days, and potentially because when the cheat is in direct competition with the producer in an environment where siderophores are less beneficial, a pleiotropic cost of the knockout is uncovered. These results therefore may provide an insight into the maintenance of cooperation in natural populations – cheat fitness is reduced by the presence of other species. Our finding is also different to that of Leinweber et al (21), who found that mixed populations of producers and cheats are competitively weaker than producer-only populations under greater cheat selection, but not under weak selection. This may be due to us using five rather than their one non-*P. aeruginosa* species, as the findings by O’Brien et al (48) showed that cheat fitness was greater when resident richness was lowest and decreased as richness increased. However, it is most likely due to us using a toxic metal to increase cheat fitness rather than iron limitation, as the former is less intraspecific and consequently has the potential to be more of a community-wide public good. This results in the cheat benefiting less from the siderophores than if they were produced for iron accumulation, and consequently they do not gain a large fitness advantage. That species can benefit from siderophore-mediated detoxification without imposing a large fitness cost to the producer has been previously found (31).

Our findings demonstrate that genotypic variation in invading populations can be highly beneficial, even when strains of a single species differ in a few genes. What is particularly interesting about this result is that the removal of these genes does not cause a direct fitness advantage to the knockout cheat strain (in fact it reduces success in monoculture). Instead, it facilitates a total overall greater invader density through equalising the fitness of the two genotypes, analogous to the genotypes filling separate niches. Whilst the importance of invader diversity for success has been previously studied, this has focused on different species of invader (49) or invaders with variation due to pre-adaption (50), with little attention given to within-species diversity of the invader. This is important to understand because natural microbial invasions are unlikely to be by populations of a single genotype, instead containing a range of genetic diversity. We do note that siderophores are important for both metal resistance and competitive ability, and genetic variation in siderophore production may have particularly large fitness effects. However, variation in siderophore production is highly relevant to natural populations, as the costs and benefits of production vary with ecological conditions meaning both producers and cheats are found in natural populations (34). Findings from previous, non-microbial, work show an increase in diversity is likely to provide an advantage to establishment probability and long-term success (49, 51, 52). Here we show this in microbes, but also that this can be context dependent.

Finding the diversity of the resident community to differ with invader type in the presence of copper but not in the absence highlights the context dependence of the impact of microbial invasions have on community composition. A surprising result here is that the most successful invasion treatment, the mixed population into the copper media, had similar diversity compared to that of the non-invaded copper media, whereas the least successful invader treatment, the cheat into copper media, caused a significant reduction in resident diversity. A likely reason for this is the significantly lower *Stenotrophomonas* population in this treatment compared to when invaded by the producer or mixed invader in the copper environment, which suggests *Stenotrophomonas* benefited from the protective effect of the invader’s siderophores. This therefore highlights that invader success and impact are not always positively associated. We do note that whilst the cheat-copper treatment had the lowest success, it was still successful, with only one out of eight replicates having a decreasing relative invader population size (i.e. with a v < 1).

Our use of a siderophore-producing invader in the context of metal pollution offers an insight into the effect of adding a metal-detoxifying microbe on the composition of a metal-polluted community. This is important to understand because metal pollution, including copper pollution, is an increasing concern globally (53). Remediating metal-polluted environments is costly and laborious, as the high persistence of metal ions means they can remain in environments for centuries. The use of siderophore-producing microbes to help remediate metal pollutants has been proposed, but it has remained unclear what the effect of their introduction will be on the resident microbial community. Here we demonstrate that when a producer strain (i.e. a detoxifying strain) is added, either in monoculture or in co-culture with the cheat, the resultant resident diversity was not significantly different between the copper-free and copper-polluted environments. This suggests that the addition of a metal detoxifying microbe into polluted environments may not significantly alter the resident diversity already present, although further work is required to test this in more environmentally relevant systems. Moreover, it suggests that this could be the case even after cheats evolve. Future work is needed to test whether these findings in our model community are extrapolatable to more natural communities and environments, and to test whether resident species will evolve to exploit the producer.

In conclusion, we show that variation in public-goods investment can provide a large fitness advantage for invading populations, but that this is dependent on the environment being invaded. This is due to the producer and cheat coexisting under higher siderophore requirement, but the producer outcompeting the cheat under weak siderophore requirement. We also show that under high siderophore requirement monocultures of the cheat have reduced success, whereas monocultures of the producer have success between that of the cheat monocultures and the cocultures. Under weak siderophore requirement, investment in production did not affect success. Finally, we show that a metal detoxifying microbe added to a polluted environment will not outcompete, and consequently reduce the diversity of, the residents in our model community. These results therefore further our understanding of the importance of cooperation for invaders, give an insight into how cooperation may be maintained, and the effect of adding a siderophore producer to a metal polluted microbial community.

## Author contributions

LL designed the study, collected the data and wrote the first draft. All authors contributed substantially to the final manuscript.

## Acknowledgments

LL and EH would like to thank the UKRI Future Leaders Fellowship award MR/V022482/1, and AB the NERC awards NE/V012347/1 and NE/S000771/1.

## Notes

### Competing Interest Statement

The authors have declared no competing interest.

